# Uncovering the Emergent Robustness and Treadmilling in a Cell-Mimicking Actin Cortex: Mechanistic insights from basic components

**DOI:** 10.1101/2024.11.04.621887

**Authors:** S Gat, Y Amit, G Livne, D. Sevilla Sanchez, N Port, A Yochelis, N.S Gov, A Bernheim-Groswasser

## Abstract

Cell shape deformation in eukaryotes, is primarily determined by the actin cortex, a thin network of actin filaments and myosin motors. This network is attached to the plasma membrane endowing cells with their mechanical stability and structural integrity. Since inside the cells hundreds of proteins are associated with the actin cortex, identifying the roles of the individual components and the mechanisms maintaining a functional cortex is highly challenging. Here, using a minimal set of components, we succeeded in recreating long-lasting dynamic treadmilling in an artificial actomyosin cortex on a bilayer membrane with cell-mimicking characteristics, both with and without myosin-II motors, replicating features observed in cells, such as myosin-induced increased cortical thickness and stress-dependent dynamics. We mechanistically reveal how treadmilling is regulated by actin network disassembly factors and myosin contractility, and the emergence of robustness of the cortical thickness to concentration fluctuations in cytoskeletal components. The robustness and responsiveness of the actin cortex which we found has a fundamental significance for the ability of Eukaryotic cells to maintain the mechanical integrity of their membrane, despite concentration fluctuation of the cytoskeleton components. Unlike complex biological cells, this system enables high-resolution and systematic studies of cortical dynamics under controlled conditions. It thus provides a cell-mimicking artificial system for designing interventions to modify the cellular cortex to improve our understanding of the basic mechanisms driving fundamental biological processes as well as for potential medical applications.

**Significance:** We report a breakthrough in understanding the actin cortex, a key structure governing cell shape, stability, and motility. By reconstructing *in-vitro* a treadmilling actomyosin cortex on a lipid bilayer using minimal components, we replicate essential cellular features. This system reveals how cortical thickness is maintained despite concentration fluctuations, highlighting an actin turnover mechanism that buffers variability. The inclusion of myosin motors is found to modify the cortical dynamics, mimicking cellular properties such as the increased thickness and stress-dependent dynamics. Unlike complex living cells, this artificial system enables systematic studies of cortical behavior, with fine control over the components and high-resolution imaging. Our artificial system provides a platform for designing interventions to modify cortical dynamics, with broad implications for biology and medicine.

**One-Sentence Summary:** In-vitro recreation of a cell-like, membrane-bound cortical actin skeleton composed of a minimal set of building-blocks with continuous turnover and emergent robustness.

## Introduction

Eukaryotic cells exhibit a variety of dynamic shapes, dependent on their function. Cell shape deformation is primarily determined by the actin cortex, a thin network of branched and unbranched actin filaments and myosin motors, that lies beneath and attached to the plasma membrane of cells *(1)*. Despite being very thin (≤ 1μm) and continuously turning over *(2, 3)*, the cell cortex endows cells with its mechanical stability and structural integrity *(4, 5)*. At the same time, its dynamic nature is essential for driving various cellular processes that involve cell shape changes such as cell division and motility *(6-8)*; some of these functions are paramount during development, immune response and cancer progression.

Two key processes govern the dynamics of the cell cortex. The first relies on the treadmilling of the actin network, which is the process of continuous polymerization at the membrane bilayer and disassembly towards the cell interior. An additional dynamical process is driven by myosin II motors which exert contractile forces within the actin cortex. All these are dissipative non-equilibrium chemical processes, involving the continuous consumption of energy (ATP) *(9, 10)*. Since hundreds of molecular components are known to interact within this cortical network *(2, 11)*, there is no understanding of the minimal set of components that are required to initiate the formation, and maintain the structural integrity of the cell cortex. The ability to control a minimal set of components is essential, for example, to designing interventions that modify the cellular cortex efficiently for medical applications.

Motivated by Feyman’s famous quote: “What I cannot make, I don’t understand”, there has been an ongoing effort to recreate the actin network under well-controlled settings using a very limited set of pure components *(12, 13)*. This approach has been previously employed to reconstitute actin cortices in various geometries *(14-18)*, but none of these previous works managed to reconstitute a membrane-bound treadmilling actin cortex, where the thickness is dynamically regulated, similar to living cells *(19-21)*.

Here, we identify for the first time, a minimal set of components and a mechanism that recreates dynamic treadmilling in an actomyosin cortex on a bilayer membrane with cell-mimicking characteristics. We demonstrate the treadmilling dynamics inside these cortices, in the absence and presence of myosin-II molecular motors. We reveal how treadmilling is regulated by actin network disassembly factors, and how this drives the robustness of the cortical thickness to concentration fluctuations in cytoskeletal components. The mechanism of the cortex emergence is formulated in a reaction-diffusion-advection model system, which explains the experimentally observed robustness. This ability of the system to efficiently buffer fluctuations in the concentration of its components through an actin turnover mechanism may be an important cellular property for maintaining proper function and responsiveness to environmental cues.

## Results

### Recreating dynamic treadmilling in an artificial actomyosin cortex on a bilayer membrane, both with and without myosin-II motors

We recreate an actin cortex assembling from the surface of a supported lipid bilayer (SLB) which dynamically turnovers for hours. Our experimental system consists of a flat PIP_2_ -based SLB-decorated flow chamber (Fig. 1A, main) and GTP-activated Cdc42 incorporated within it, nucleation promoting factors (NPF) (specifically, ΔEVH1 N-WASP, the near full-length version of N-WASP lacking only the N-terminal EVH1 domain *(22)*, which localize to this SLB and physically link the actin cortex to it (Fig. 1A). The system also includes a solution of a well-defined and controlled set of purified components which control the rate of actin assembly/disassembly and can also be supplemented with myosin-II motor filaments (Fig. 1A). Using intrinsically inactive N-WASP, instead of its intrinsically active VCA form *(23)*, not only better represents the native biological system, but also ensures that actin network assembly is exclusively on the membrane surface (Fig. S1), where N-WASP is activated by binding to both PIP_2_ and Cdc42 *(24)*. In particular, the use of a flat geometry enables to quantitatively characterize the intrinsic dynamics of actin cortex growth and turnover, decoupled from curvature effects *(25)*. The chamber volume to surface area and the employed concentrations have been purposely designed to mimic the restricted availability of components observed in cellular environments: In our chamber the ratio of available actin, the most abundant protein in most eukaryotic cells *(26, 27)*, to membrane area (where actin network assembly takes place), is comparable to the value for typical cells (see SI).

**Fig. 1.**
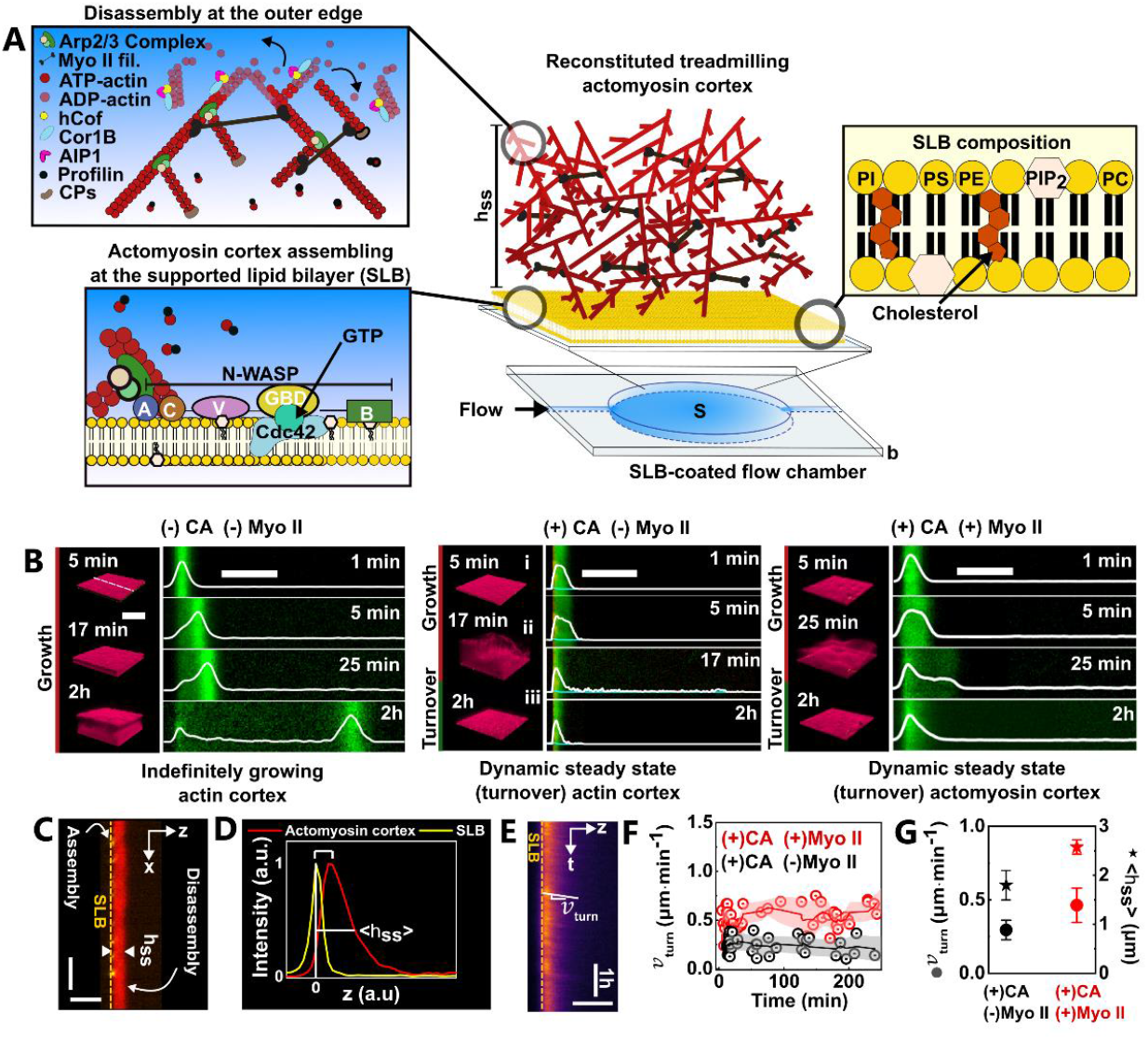
Reconstitution of a long-lasting dynamic steady-state actin cortex on a supported lipid bilayer (SLB). (**A**) Schematic of the experimental setup: a flow chamber with a supported lipid bilayer (SLB, yellow), actin nucleation factors which trigger actin cortex assembly and accessory proteins regulating actin assembly/disassembly dynamics, and myosin II motors. *h*_*ss*_ is the steady-state thickness. S and b are the chamber surface and thickness, respectively. (**B**) Sequences of 3D (actin - red) and side-view (*xz* cross-section measured along the dashed white line) (actin - green) confocal images showing the spatio-temporal evolution of actin cortices assembling from an SLB (yellow), with and without disassembly factors and/or myosin motors. Shown are actin cortices that either continuously grow with time (left panel) or evolve continuously into a dynamic turnover steady state (middle and right panels). White curves depict the average intensity distribution of F-actin measured normal to the SLB (*N* = 115). (**C**) Side view confocal image of an SLB-bound actomyosin cortex of thickness *h*_*ss*_ at steady state. (**D**)Spatial distribution of the actin network (red) and membrane (yellow) measured perpendicular to the coverslip. Actin density decays exponentially and its peak position is *z*-shifted with respect to the SLB; the steady state thickness is measured at half-maximal intensity. (**E**)Turnover speed (*v*_*turn*_) extracted from the slope of Kymograph plots, showing steady actin fluxes and turnover velocity over time. (**F**) Turnover speed for long-lasting treadmilling actin cortices with and without myosin motors. (**G**) Turnover speed and steady state thickness (mean ± SD, *N* = 40 − 50). Conditions: 25 nM CA (B middle and right panels, C, E, F, and G), 50 nM Myosin II (B right panel, C, E, F, and G). Scales bars are 10μm.

Aiming to identify what constitutes a minimal set of pure components that allows for a treadmilling actin cortex of finite steady state thickness *h*_*ss*_ to form(Fig. 1A), we include actin monomers along with accessory proteins such as Profilin, Capping proteins (CPs), and Arp2/3 complex, to regulate the rates of actin cortex assembly. Their concentration is kept constant in all studied systems. Network disassembly rate is regulated by a mixture of three components (Cor1B, AIP1, and hCof), which facilitate the disassembly, debranching and severing of ADP-bound actin filaments (F-actin) (Fig. 1A) *(28-31)*. We maintain a constant concentration of hCof, and modulate the disassembly rates by adjusting the concentrations of Cor1B (C) and AIP1(A) between 0 − 75 nM, keeping a 1: 1 molar ratio (C: A) *(29)*. Additionally, in some of the experiments we have added 50nM myosin II motors in the form of small filaments (or aggregates) of controlled size *(32, 33)*, roughly consisting of 50 myosin dimers, similar to their size in cells *(34)*. All experiments are performed with 2mM ATP, maintained constant using an ATP regenerating system, which keeps actin and myosin-dependent activities steady for several hours *(32, 35-37)*.

Initially (at *t* = 0), we introduce the components into the chamber. This prompts the rapid (less than 1min) formation of a thin and tightly packed cortical layer on the SLB surface (Fig.1B). This thin initial cortical actin network forms with very similar thickness (denoted as *h*_0_, see Fig. 2A) and density profile for all studied systems (Fig. 1B). The progression towards steady state involves a transient, nonlinear ‘growth’ phase during which the network thickens over time (Figs. 1B and 2A) while its density decreases (Fig. 1B, side view images). Network thickness growth can be accompanied by large thickness fluctuations (Figs. 1B middle and right panels and 2A (main panel)), which may indicate that non-uniform stresses develop in the network. The gradual density decrease occurs throughout the thickness and is most clearly visible at the cortex’s outer edge, where the network is oldest (see density profile variations in Fig. 1B). It mirrors the progressive disassembly of the actin network with time, clearly more efficient in the presence of CA (Fig. 1B, middle and right panels at times 1 − 17min and 1 − 25min, respectively). This process replenishes the system with components (such as actin monomers, Arp2/3 etc.) that diffuse to and repolymerize at the SLB surface, promoting the fresh formation of an actin network (see the density profile variations in Fig. 1B). The actin growth at the SLB pushes the preformed network upwards, leading to the observed thickening effect. This transient growth regime culminates in a rapid thickness reduction following disassembly of the cortex’s outer edge (Figs.1B middle and right panels, 2A - (ii) main panel, and Movie S1). Note that systems lacking CA remain in the ‘growth’ phase over the few hours timescale of the experiment (Figs. 1B left panel and 2A (inset) depict the first two hours out of six, Movie S2), highlighting the pivotal role of network disassembly in achieving a dynamic steady-state.

**Fig. 2.**
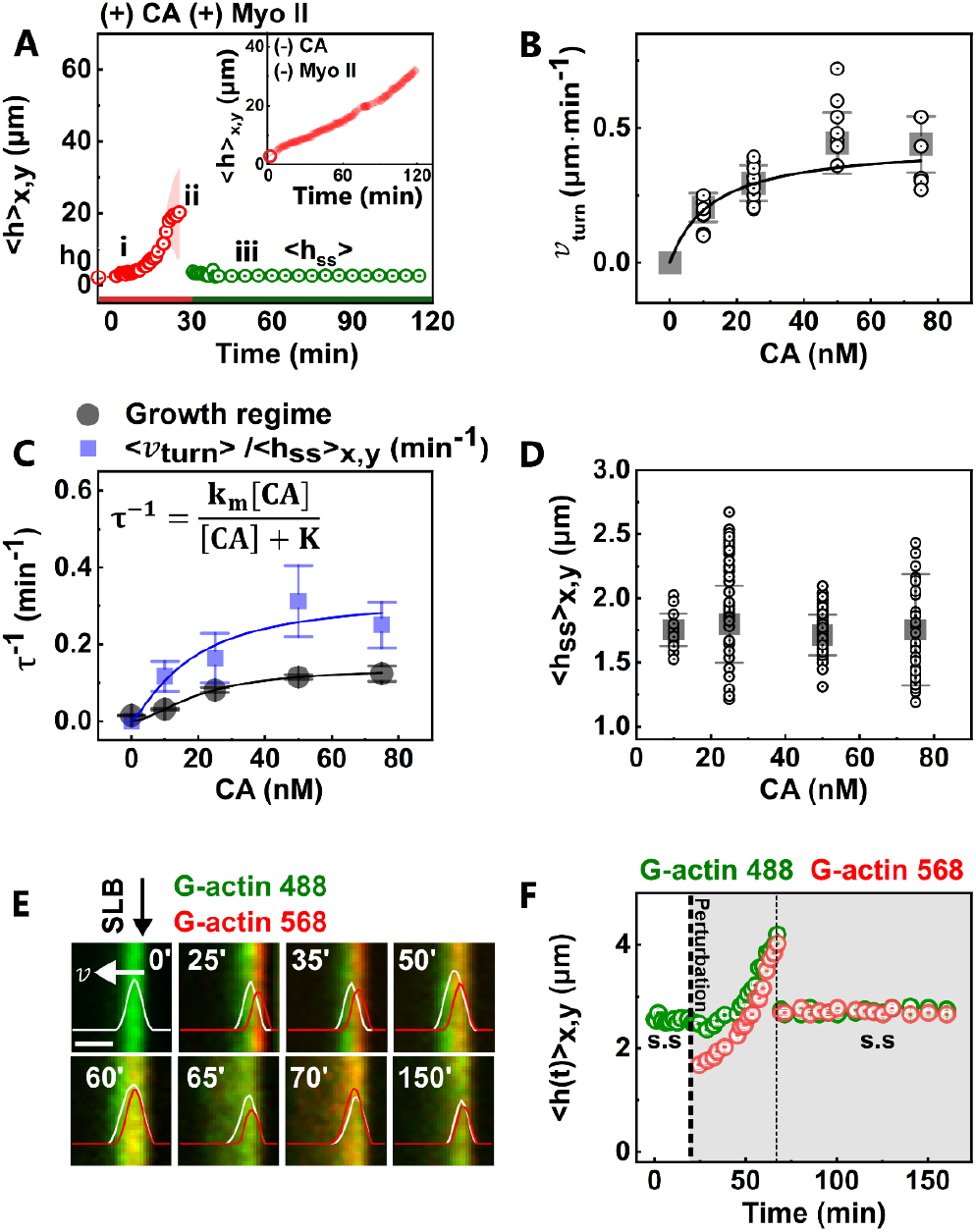
Actin cortex turnover speed and steady-state thickness dependence on disassembly factors in absence of myosin. (**A**) (main) Cortex thickness over time for the cortical networks in Fig. 1B right (main) and left (inset) panels with individual data (mean) and ribbons (SD). The growth (i, ii) and steady state (iii) regimes after 5, 11, and 60 minutes. *h*_0_ denotes the initial cortical actin network thickness. (**B**) Turnover velocity *v*_*turn*_ vs. [CA], with individual data points and mean ± SD (*N* = 40 − 50 per [CA]), and a fit line (*r*^2^ = 0.9*8*). (**C**) Actin disassembly rate (**τ**^−1^) vs. [CA], determined from actin intensity decay during the transient growth phase (black) and from the turnover velocity-to-thickness ratio in the steady-state regime (blue), with fit lines (*r*^2^ = 0.95). (**D**) Surface averaged steady-state cortex thickness (< *h*_*ss*_ > _*x,y*_) vs. [CA], with individual data points and mean ± SD (*N* = 100), and a fit line (*r*^2^ = 0.96). (**E-F**) Response to an external perturbation: actin cortex at steady state (green) is supplemented with 2µM red actin monomers and accessory proteins at *t* = 23 min, showing incorporation, growth (*v* marks the growth speed and the arrow its direction), and disassembly. Average steady state thickness changes from 2.54 ± 0.05 μm (before perturbation) to 2.72 ± 0.05 μm (after perturbation), corresponding to a 10% increase. (**E**) Bar is 5μm and [CA] = 60nM (prior and after perturbation).

After this transient stage, and only in the presence of CA (with or without myosin II), the system stabilizes at its steady state thickness, *h*_*ss*_ (Fig. 1C) which remains stable for hours (Figs. 1B and 2A - (iii) main panel depict the first two hours out of five). These systems exhibit a characteristic asymmetric actin density distribution across their thickness (Figs. 1B middle and left panels, 1D, and S2), similar for all studied systems, with the peak density spatially shifted relative to that of the membrane (set at *z* = 0) and decaying exponentially away from the SLB surface (Figs. 1D and S2), mirroring observations in living cells *(19, 38)*.

While all systems containing a finite amount of CA reach such long-lasting (i.e., hours) treadmilling steady-states, the transition time *t*_*g*→*ss*_ from growth to steady state regime gets shorter with the increase in [CA] due to faster disassembly rates, reducing to mere minutes at elevated CA levels (Fig. S3). Intriguingly, the presence of myosin II delays the arrival to a steady state compared to systems lacking motors (Figs. 1B and S3), possibly due to competition with disassembly factors for F-actin binding or due to internally applied active stresses generated by myosin that may slow down the network disassembly during the transient phase before the onset of the steady-state turnover.

### The steady state regime, with and without myosin, is long-lasting and characterized by a continuous upward actin flux

We turn now to characterize the steady state regime. We find that it is characterized by a continuous upward actin flux starting from the SLB surface (Fig.1E, F, and Movie S3). The actin turnover velocity, *v*_*turn*_, is extracted from the slopes of kymograph plots (Fig. 1E). These trajectories are straight, suggesting that the actin speed remains uniform across the thickness – a clear sign that the cortical network is moving away from the membrane as a solid, in a dynamic steady state. Over the entire course of the experiment new trajectories consistently emerge, all displaying similar linear slopes, indicating that both the actin fluxes and the turnover velocity are long-lasting and stable over time (Fig. 1F). The steady-state turnover velocity is stable both in the presence and absence of molecular motors, but it is significantly higher when myosin-II is present (Figs. 1F and G), which is also the case for the steady state thickness (Fig. 1G)

### Dependence of the turnover velocity, cortex thickness, and network disassembly rate on disassembly factors (CA) concentration

In Fig. 2B we plot the dependence of *v*_*turn*_ on CA concentration, observing a nonlinear monotonous increase up to saturation. Similarly, we plot the network disassembly rate **τ**^−1^ in Fig. 2C. This quantity is only accessible during the transient ‘growth’ phase, when we can evaluate it from the measured decay of the fluorescence of the F-actin density at the cortex’s outer edge over time (Figs. 1B, S4, and SI Materials and Methods). We find that the average F-actin density at steady state decays exponentially in space (Fig. S2) and with time (Fig. S4), the latter implies a roughly constant disassembly rate. In Fig. 2D we plot the average steady-state thickness of the actin cortex and find that unlike *v*_*turn*_ and **τ**^−1^, it is roughly independent of the CA concentration.

To gain mechanistic insights into the system behavior, we construct a one-dimensional reaction-diffusion-advection model, accounting for concentrations of F-actin (*p*), G-actin (*m*), freely diffusing disassembly agents (*c*_*f*_), and F-actin decorated by disassembly agents (*c*_*b*_). The model is described schematically in Fig. 3A, while the equations are given in the SI. Note that the model focuses on the dynamics of the disassembly factors *(39)* and does not include myosin motors *(40, 41)*. In Figs. 3B,C we show that the numerical solutions indeed reproduce the experimentally observed transients and the steady-state characteristics, such as the exponential decay of the F-actin (i.e., *p* + *c*_*b*_) density in space, insensitivity of ⟨*h*_*ss*_ ⟩ to the CA concentration (i.e., *c*_*f*_), and the nonlinear dependence of the turnover speed *v*_*turn*_ on the CA concentration. Both experiments and the model solutions capture the fast initial formation of a thin polymerized layer, which then thickens during the growth phase (with a local high-density region that is pushed away while decaying in amplitude), and the final formation of a thinner steady-state exponential profile (Fig. 3B).

**Fig. 3.**
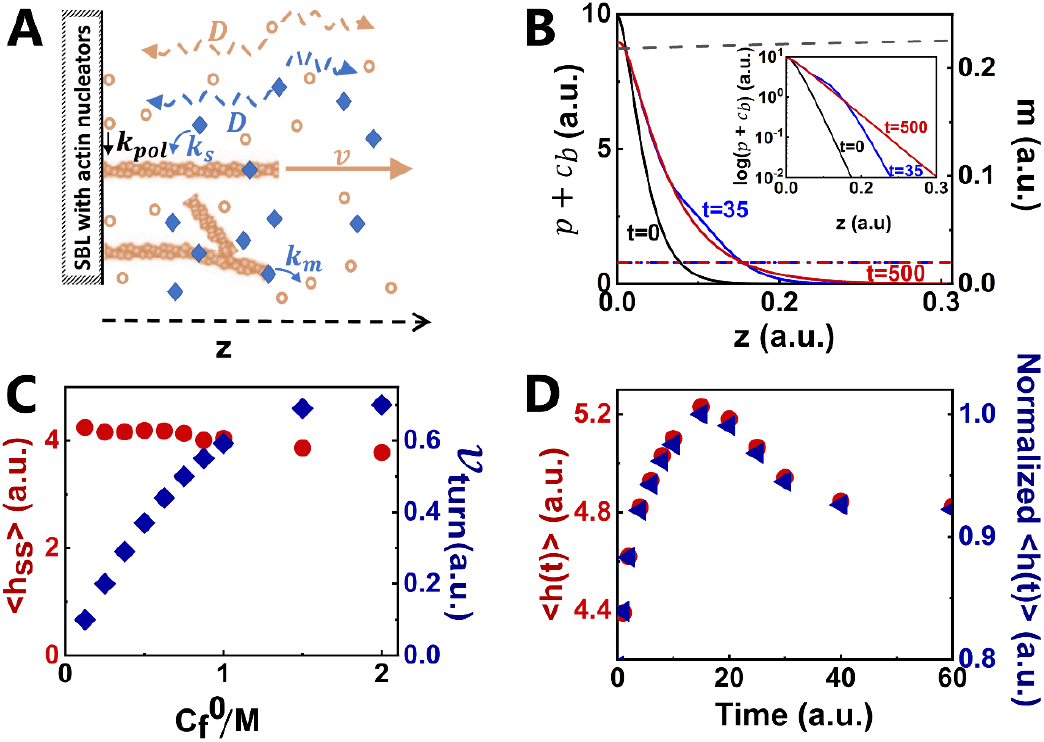
Reaction-diffusion-advection model system describing the mechanism of cortex emergence. (**A**) Schematic description of the treadmilling cortical network model (see SI for details). (**B**) Model-calculated spatial profiles at different times (solid lines), with dashed lines representing actin monomer distribution. (**C**) Steady-state thickness (red dots) and turnover velocity (blue diamonds) for varying disassembly factor concentrations relative to actin monomers, with thickness measured at half-maximal polymerized actin distribution. (**D**) Simulated width of polymerized actin distribution vs. time following actin monomer addition, showing a 10% increase in cortex thickness.

Next, we show that the steady state properties are in fact related to the disassembly rate and the rate of actin turnover at the SLB. The disassembly occurs at an effective rate **τ**^−1^ and in our model arises from two sequential first order reactions: (i) free disassembly factors bind to newly polymerized actin at rate *k*_*s*_ per [CA] and (ii) the actin filaments decorated by disassembly factors (i.e., *c*_*b*_) undergo disassembly (with rate *k*_*m*_), releasing actin monomers and free disassembly factors into the solution. These two processes give rise to the following effective rate at which polymerized actin is disassembled, given by (see SI Eq. S7*(42)*):

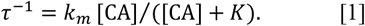

In Fig. 2C (black dots) we fit the experimental data to Eq. 1, extracting the values of *K* = *k*_*m*_ /*k*_*s*_ = 19.7 ± 2.6 nM and *k*_*m*_ = 0.13 ± 0.01 *s*^−1^. In the regime of fast diffusive transport of free monomers (*m*) from the bulk to the SLB, the polymerization rate that governs the cortical advection speed is in turn dependent on the supply of monomers, which is dictated by the disassembly rate. We therefore expect that

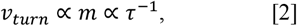

see also the detailed model equations and the choice of model parameters in the SI. Consequently, by fitting the experimental data of *v*_*turn*_ in Fig. 2B to the functional form of Eq. 1 we obtain *K* = 19.6 ± 5.5nM which agrees with the dependence found in Fig. 2C and thus, validates Eq. 2. From the reaction-diffusion-advection model in the SI, we describe a polymerized actin cortex that is pushed from the SLB at speed *v*_*turn*_, and undergoes disassembly at the effective rate given in Eq. 1. This means that at steady state the profile of the actin cortex is expected to be exponential (see SI Eqs. S9, S10, and inset of Fig. 3B) with thickness given by the ratio: *h*_*ss*_ = *v*_*turn*_ /**τ**^−1^. From the discussion above and from Eq. 2, we therefore expect this steady-state thickness to be independent of the CA concentration, as we find in the experiments (Fig. 2D).

To compare quantitatively, we note that the **τ**^−1^ cannot be measured during the steady state regime, only during the initial transient growth phase (Fig. S4). Nevertheless, we can estimate it from the ratio *v*_*turn*_ /*h*_*ss*_ measured at steady state. In Fig. 2C (blue dots) we see that this calculated ratio is ∼2.5 times higher than the measured **τ**^−1^ during the initial growth phase, indicating that the disassembly rate at the transient regime is slower by this factor compared to the disassembly rate at the steady state. This difference could arise from the higher network density and rigidity of the thin initial cortical actin layer compared to the steady state cortex. Note that the functional dependence of **τ**^−1^ on the CA concentration is the same during the growth and steady state, which indicates that *K* is a constant in Eq.1. This therefore means that while the overall rate is different (due to larger *k*_*m*_ = 0.31 ± 0.03 *s*^−1^ in the steady state regime), the ratio 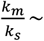 *const*. The origin of this behavior could be that both the binding of disassembly factors and their ability to disassemble depend in a similar manner on the F-actin age and network density (see SI).

### Response of the system to an external perturbation shows robustness of the cortical thickness to concentration fluctuations in cytoskeletal components

After showing that the system sustains a dynamic steady-state with continuous turnover, we wanted to explore its response to an external perturbation. In Fig. 2E we show a steady-state cortex to which we rapidly added an additional supply of 2µM Alexa 568-actin monomers together with accessory proteins kept at their original concentration (prior to the perturbation). The figure clearly shows that the monomers are incorporated into the growing F-actin cortex at the SLB, and then advected and degraded. Following the additional supply of monomers at the SLB, the cortex thickness increases (Fig. 2F) but later returns to the steady-state behavior with a thickness that is increased by the proportion of added actin (by about ∼10%). We can simulate this experiment using our 1D model, by similarly adding uniformly a small amount of actin monomers to the system which was already in a steady state.

Since the concentration of freely diffusing disassembly agents (*c*_*f*_) has a negligible effect on the cortex thickness (Fig. 3C), in the simulation we only added G-actin monomers (*m*). As observed in the experiments, we find that the model predicts a transient increase in the cortex thickness, before it returns to a new, slightly thicker, steady state (Fig. 3D). This agreement further supports our advection-degradation model of the turnover and advection dynamics of the actin cortex.

### Effects of myosin II on actin cortex dynamics reveals a myosin-induced increased cortical thickness and stress-dependent dynamics

We now turn to exploring the effects of myosin II on actin cortex dynamics, as shown in Fig. 4. In the absence of CA the cortex continues to grow (Fig. 4A, C), similar to the system in the absence of motors (Fig. 1B left panel), but with a slower rate (inset of Fig. 4C). This significant slowing down of the thickness growth in the presence of the myosin motors could arise from active forces that contract the actin network as well as the motors slowing down the residual disassembly rate. In the presence of CA, the initial growth phase is later replaced by a thin cortical layer at the steady-state conditions (Fig. 4B and F (inset)), similar to the behavior in the absence of motors (Fig. 1B middle panel).

**Fig. 4.**
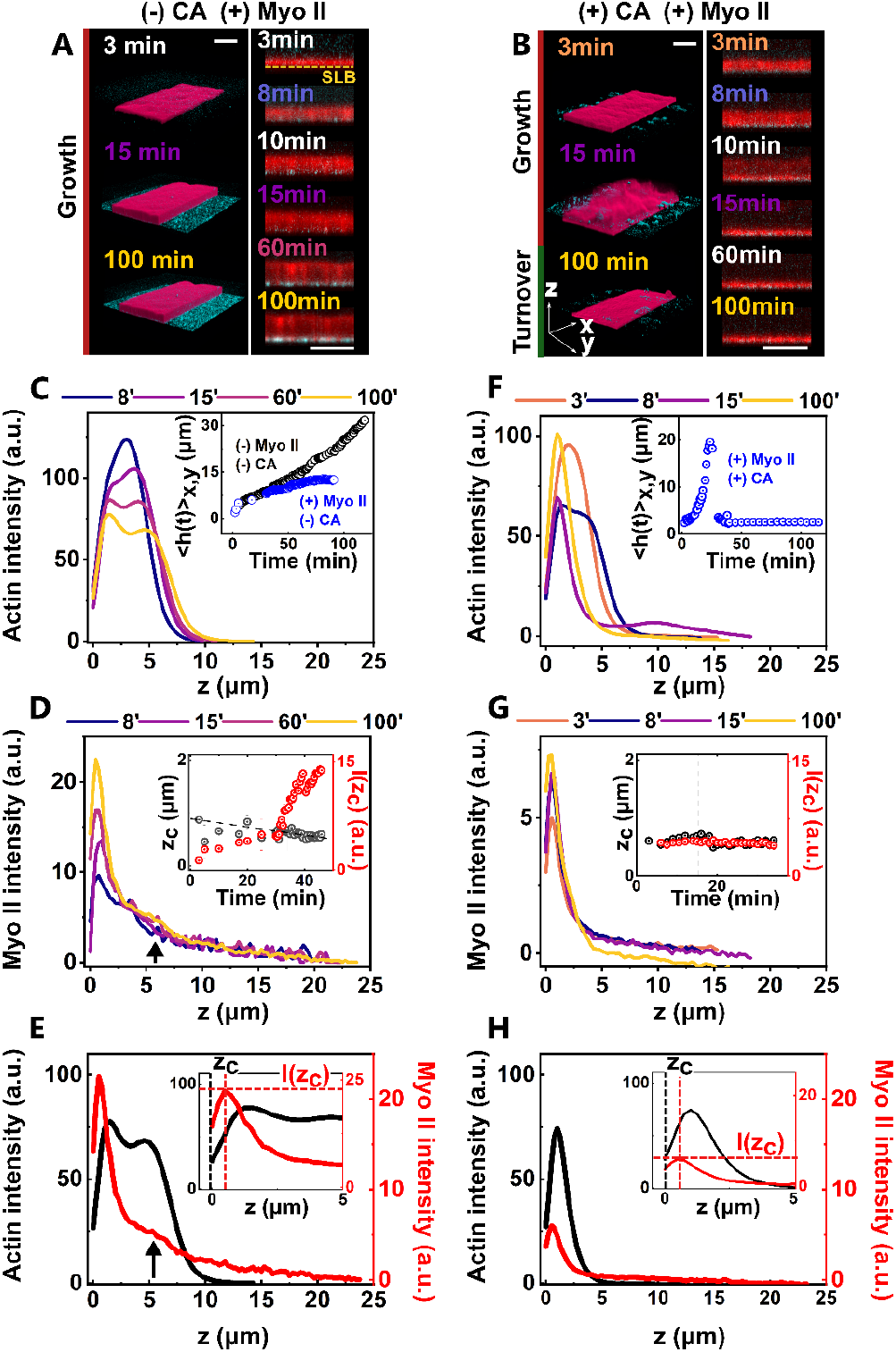
Actin turnover and myosin contractility govern cortex growth and turnover dynamics. (**A-B**) 3D and side-view confocal images showing the spatio-temporal evolution of actin cortices (red) assembling from an SLB surface (not shown) with myosin (cyan), without (A) and with (B) disassembly factors CA. (**C-E**) Effect of myosin contractility on actin cortex dynamics without disassembly factors. (**C**) Actin and (**D**) myosin density profiles measured normal to the SLB at different times. (**E**) Combined density distributions at *t* = 100 min, with the red dashed lines indicating the average myosin density peak (*z*_c_) and intensity (*I*(*z*_c_)), which correlates with actin density at the cortex rear. Insets: show average cortex thickness vs. time, with (blue) and without (black) myosin (**C**) and zoom-in of the temporal evolution of *z*_c_ and *I*(*z*_c_) (**E**). (**F-H**) Same as (**C-E**) with 25nM CA. Myosin concentration is 50nM. Scales bars are 10μm.

Both with and without CA, we find that the myosin motors (Fig. 4A, B cyan) accumulate over time at the SLB, which is consistent with their plus-end persistent movement which directs them to the growing tips of the actin network (Fig. 4A-H, Movie S4). In the absence of CA (Fig. 4D, E) the accumulation of motors at the SLB reaches much higher values compared to the system with CA (Fig. 4G, H, and Movie S5). This may be attributed to the much thicker actin layer when CA is absent, which adsorbs more motors from the solution (“antenna-effect” *(43)*). The continuous growth in myosin intensity at the SLB also indicates that this system does not reach a real steady state. We note also that the myosin distribution forms an additional small accumulation which follows the second peak of the actin distribution (see arrow in Fig. 4D, E). The roughly constant density, step-like actin cortex observed in the absence of CA can be attributed to the contractile forces of myosin motors exerted within the network (compare to the absence of motors in Fig. 1B left), as predicted in *(40)*.

However, in the presence of CA and actin treadmilling (Fig. 1E, F), the myosin intensity reaches rapidly a constant steady-state value (inset of Fig. 4G), even faster than the actin distribution which exhibits a transient growth phase (Fig. 4F). In the presence of high turnover rates, the actin distribution is exponential and its functional form is not affected by the motors (in agreement with *(40)*). Note that both in the presence and absence of CA we find that the long-time myosin distribution has a very broad distribution that extends well beyond the visible polymerized actin layer (Fig. 4E,H). In addition, we cannot visualize the motion of individual myosin filaments, but occasionally can track the motion of large aggregates of motors which are observed to move bidirectionally in this network, some move towards the SLB and accumulate there, while others are pushed by the actin treadmilling to the edge of the cortex, where they can be released into the solution (Fig. S5A and Movies S5, S6). We note that the steady state shown above resides in an intermediate range of values of the concentration of the disassembly factors (Fig. 5). Below this range the actin cortex seems to be “jammed” and no turnover is observed (Fig. S5B), while above this range the cortex thickness does not relax to a constant value and undergoes repeated bursts of growth and decay (Fig. S5C and Movie S6).

**Fig. 5.**
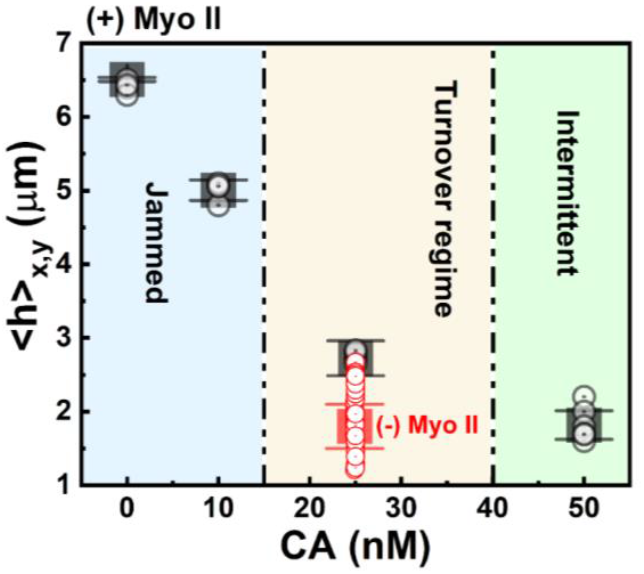
Combined effects of myosin contractility and actin disassembly on the dynamical regimes and average cortex thickness at long times (*t* = 60min). Shown are individual data points and mean ± SD (*N* = 110) for 50nM myosin II (black) and without myosin II (red).

Surprisingly, the steady state cortex with myosin motors is significantly thicker compared to the cortex without motors (Figs. 1G and 5), which corresponds to a significant increase in the turnover speed (Fig. 1F, G). By taking the ratio of the turnover speed to the thickness we can estimate the disassembly rate, which we find to be only slightly enhanced by the presence of myosin (**τ**^−1^ = 0.19*8* ± 0.03 *s*^−1^ and 0.164 ± 0.03 *s*^−1^, with and without myosin), possibly due to the low myosin concentration (at high concentration myosin can promote significant actin disassembly *(44, 45)*). This indicates that the thicker cortical layer in the presence of motors is primarily due to the increase in actin polymerization speed. This effect of the myosin motors could arise from the motors locally pulling the network, as well as bending and deforming the actin filaments at the SLB such that there is more available space for the incorporation of new monomers. This observation is in qualitative agreement with measurements of the actin cortex thickness in living cells, which decreases when myosin motor activity is inhibited *(21)*.

## Conclusion

In the present work, we report, for the first time, an *in-vitro* system of minimal number of purified components that exhibits long-lasting steady-state treadmilling actin cortex on a supported cell-mimicking lipid bilayer (SLB). Within these artificial membrane-bound cortices, the actin turnover is polarized with polymerization confined to the membrane, depolymerization occurring away from the membrane, creating a detectable treadmilling flow that is perpendicular to the membrane. While we focused on a particular set of purified components, there may be other minimal sets that could support treadmilling conditions, due to the redundancy within cells. We suggest a generic theoretical mechanism which shows qualitative agreements not only with steady-state conditions but also with transient dynamics.

We find that the treadmilling actin cortex exhibits emergent robustness of its thickness to variations in the concentration of the disassembly components that facilitate the turnover flow. Using a mathematical model, we identified the mechanism which drives this robustness, namely that the actin disassembly rate determines also the rate of cortical flow through the supply of actin monomers. This suggests that cells can utilize this mechanism to buffer concentration fluctuations and maintain a functional cortex of sufficient thickness, which is insensitive to local variations in the concentration of the disassembly machinery. We also show that transient fluctuations in the actin concentrations cause thickness variations, which quickly decay away in this dynamical system. Further studies of the cortice’s robustness and response to different perturbations can now be carried out using this minimal system.

The myosin motors in our experiments get embedded into the artificial actin cortex, accumulating at the growing tips, as observed in cells in a cell-cycle dependent manner *(38)*. The presence of myosin motors produced a thicker cortical network, which we attribute to the significant myosin dependent increase in the rate of actin treadmilling speed. This feature has not been directly observed previously and suggests that future theoretical models of this system should include stress-dependent biochemical reaction rates *(25, 36, 46, 47)*. The thickness increase in the presence of myosin motors agrees with observations of cortical actin thickness in living cells *(21)*. However, in cells the myosin activity also induced large and localized thickness fluctuations *(41)*, which we do not observe here. The intermittent regime of repeating cycles of cortex growth and dissipation which we observed is more similar to the periodic cycles of cortex growth, condensation and detachment observed in cell extracts *(48)* and motile cell fragments *(49)*.

Our model system has an open configuration that allows the study of the actin cortex dynamics with very high spatio-temporal resolution, and highly controlled temporal manipulations of the composition, which are difficult to achieve in living cells. Consequently, the proposed system paves the way for future systematic studies of cell-like cortical dynamics. Moreover, it should also allow for systematic study of new medical interventions, such as facilitating drug design and testing in a cell-mimicking artificial system. We, therefore, believe that this work provides a fundamental leap to our understanding of the basic mechanisms driving fundamental biological processes as well as potential medical applications.

## Materials and methods summary

### Materials

All lipids are purchased from Avanti Polar Lipids except for the labeled PIP_2_ that is purchased from Echelon Biosciences. GTPγs is purchased from ROCHE and the Arp2/3 complex from Cytoskeleton Inc.

### Protein purification

G-actin is purified from rabbit skeletal muscle acetone powder by gel filtration *(50)*, stored on ice, and used within 3-4 weeks. Actin is labeled on Cys374 with Alexa-Fluor 488 C_5_ maleimide or Alexa-Fluor 568 C_5_ maleimide (Invitrogen). Myosin II skeletal muscle is purified following *(51)* and labeled with Alexa-Fluor 568 at pairs of engineered cysteine residues *(52)*. The other proteins used in this study are described in the SI.

### Preparation of dynamic steady state actin cortex on supported lipid membranes

Flat SLBs are prepared by incubating the chamber with a solution of 1mg/mL small unilamellar vesicles (SUVs) of cell-mimicking lipid composition (mol %: 30% DOPE, 30% DOPC, 10% liver PI, 25% DOPS, 4.9% Brain PIP_2_, 0.1% BODIPY TMR-PIP_2_, and a supplement of 15% cholesterol) for 15min at RT. The chamber is then washed and incubated with 2μM GTPγs-activated Cdc42 for 7min at 37°C, followed by the incubation of 1μM ΔEVH1 N-WASP for 15 min at RT. The chamber is washed and then loaded with the actin solution which sets the experiment’s starting time. In the experiments we use an actin solution that contains a fixed amount of actin monomers (10μM), Arp2/3 complex (40nM), Profilin (2.5μM), CPs (50nM), and hCof (3μM). It is supplemented with variable amounts of Cor1B and AIP1 (at 1:1 molar ratio), which can vary between 0 − 75 nM. Myosin II motors are added in the form of small filaments, roughly consisting of 50 myosin dimers *(32)*. The solution also includes 2mM ATP, maintained constant using a, Creatin Kinase and Creatin Phosphate, ATP-regenerating system *(32)*. The molar percentage of labeled actin monomers and myosin motors is 10 and 50 %, unless stated otherwise.

### Microscopy techniques

Samples are imaged using a laser scanning confocal microscope (LSM *88*0, Zeiss Germany) in fast Airyscan mode and controlled via the ZEN software (Zeiss Germany) or with a 3i Mariana (Denver, CO) spinning disk confocal microscope, equipped with a Yokogawa W1 module and a Prime 95B sCMOS camera, and controlled by Slide Book software. Actin is excited at 4*88* nm and, PIP_2_ or myosin motors, at 56*8*nm.

### Data quantification

The raw images are corrected for photobleaching, and for lateral and axial drifts using a ‘registration’ function from the ANTs package *(53)*. We employ a Rigid transformation algorithm (which includes both rotation and translation) to correct the drift within the *x,y* plane and *z*-directions, passing Dense sampling parameters for achieving higher accuracy. From the stabilized confocal images, we extract the mean intensity of the actin monomers and myosin motors in the bulk solution, from which we deduce the densities of the polymerized (F-actin) network and the myosin II motors and their distributions across the cortex thickness. For data quantification we employ image segmentation to define the actin cortex, top and bottom, surfaces, which we combine with the normal density distributions of the F-actin, myosin, and the SLB to characterize the actin cortex thickness dynamics (spatio-temporal evolution and steady state properties), F-actin disassembly rate **τ**^−1^, and actin network turnover speed, *v*_turn_. Moreover, we characterize the active transport of individual large motor clusters in the network and the dynamics of the smaller myosin II filaments’ population.

Data quantification is performed using MATLAB (MathWorks, MA, USA), Python, Origin (OriginLab Corp., MA, USA), and ImageJ software. For 3D image reconstruction and data deconvolution we use the Huygens Professional 21.04 software package (Huygens; Scientific Volume Imaging, Hilversum, the Netherlands).

Due to space limitation a comprehensive Method section is provided in the SI (Material and Methods section).

## Acknowledgments

We thank Karsten Kruse, Itamar Kolvin, Marco Fritzsche, Maayan Levi Segoli, and Uzi Hadad for useful discussions. We thank Shira Albeck and Tamar Unger for proteins expression and purification at the “Israel Structural Proteomics Center (ISPC)” at the Weizmann institute of science. We thank Dina Aranovich for protein purification and labelling. We thank Pekka Lappalainen for providing the CPs construct, Henry Higgs for the Profilin constructs, Laurent Blanchoin for the hCof construct, Bruce L. Goode for the Cor1B and AIP1 constructs, Wendell A. Lim for the ΔEVH1 N-WASP construct, and Richard A. Cerione for the Human Cdc42 construct.

## Author contributions

SG Designed and performed experiments, developed all analytical methods for data quantification, analyzed the experimental results, prepared the Figures and Movies, and wrote the manuscript. YA Developed analytical methods for data quantification. GL Developed analytical methods for data quantification. DSS Developed analytical methods for data quantification. NP Wrote the mathematical model. AY Wrote the mathematical model and the manuscript, performed the numerical computations. NSG Wrote the mathematical model and the manuscript. ABG Designed and developed the experimental system, developed analytical methods for data quantification, analyzed the experimental results, and wrote the manuscript.

## Data availability

All data are available in the manuscript or the supplementary materials.

## Supplementary Materials

### Materials and Methods

#### Materials

Arp2/3 complex is purchased from Cytoskeleton Inc (Denver, USA). Non-hydrolysable GTP, Guanosine 5′-[γ-thio]triphosphate tetralithium salt (GTPγs, CAS no. 94825-44-2) is purchased from ROCHE. Lipids are from Avanti polar lipids and purchased in their solubilized form (in chloroform, CHCl_3_): 1,2-dioleyol-sn-glycero-3-phosphaethanolamine (DOPE cat# 850725C), 1,2-dioleoyel-sn-glycero-3-phosphacholine (DOPC cat# 850375C), L-α-phosphatidylinositol (Liver PI cat# 840042C), and 1,2-dioleoyl-sn-Glycero-3-Phospa–L-Serine (DOPS cat# 840035C). Owing to the high negative charge of phosphatidylinositol-4,5-bisphosphate (PIP_2_) it is solubilized in CHCl3:Methanol:H2O 20:9:1 (v/v/v) (PtdIns(4,5)P_2_ cat# 840046X). Purchased as powders are: Cholesterol (cat# 700000P) from Avanti Polar Lipids and BODIPY TMR-PIP_2_ (cat# C-45M16a, Echelon Biosciences). Both are solubilized in chloroform (cat# 308022100), kept under Argon atmosphere and protected from light, like all other lipid solutions. Lipid mixtures are prepared by mixing appropriate volumes of the lipid and cholesterol stock solutions. The CHCl_3_ is then evaporated for 3 − 4 h at under vacuum (50 mbar) in a rotating evaporator (V-800, BÜCHI). The dried lipid mixture is kept under Argon atmosphere at −*8*0°C until used.

#### Methods

##### Experiments design and experimental procedures

###### Proteins purification

G-actin is purified from rabbit skeletal muscle acetone powder by gel filtration *(1)* (HiPrep™ 26/60 Sephacryl™ S-300HR, GE Healthcare), stored on ice, and used within 4 weeks. Actin is labeled on Cys374 with Alexa-488 C_5_ maleimide or Alexa-568 C_5_ maleimide. Purification of myosin II skeletal muscle is performed following standard protocols *(2)*. Myosin II is labeled with Alexa-568 C_5_ maleimide at pairs of engineered cysteine residues as in *(3)*. The construct of recombinant mouse CPs is a kind gift of Pekka Lappalainen and purified following *(4)*. Profilin construct is kindly provided by Henry Higgs and purified following *(5)*. The construct of recombinant human cofilin (hCof) is a kind gift of Laurent Blanchoin and purified following *(6)*. Cor1B and AIP1 mouse constructs are kindly provided by Bruce L. Goode and purified according to *(7)*. N-WASP ΔEVH1, the near full-length version of N-WASP lacking only the N-terminal EVH1 domain, construct is kindly provided by Wendell A. Lim and purified following *(8)*. Human Cdc42 construct is kindly provided by Richard A. Cerione and purified as a His6-tag protein by baculovirus-mediated expression in (Sf9) insect cells following *(9)*. The concentration of all proteins is determined by absorbance using theoretical extinction coefficients.

###### Flow chamber design

The flow chamber is intentionally designed to, along with the concentration of employed pure components, mimic the restricted availability of components observed in cellular environments. Our flow chamber has a circular shape (disc area S = 230mm^2^) and is assembled from two glass coverslips spaced b = 140μm apart, such that the ratio of the chamber volume *V* = S ·b to the SLB total area (2S) is constant b/2 = 70μm, and independent of the SLB area.

*Cortex assembly dynamics under restricted components availability*: we estimate the limited availability of components in our system from the ratio of available moles of actin, the most abundant protein in most eukaryotic cells *(10, 11)* to the total SLB area (where actin network assembly takes place):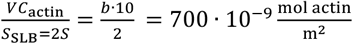, where *C*_actin_ = 10μM in our experiments. This value is comparable to that for typical eukaryotic cells: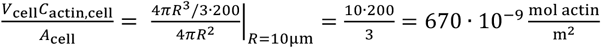, where *V*_cell_ and *A*_cell_ are the volume and surface area of the cell, R = 10 μm is a typical cell radius, and *C*_actin,cell_ = 200μM is the typical cellular concentration of actin *(10, 11)*.

###### Minimal systems that exhibit cortical dynamic turnover on suspended lipid bilayers (SLBs) – experimental procedure

Flat SLBs are prepared by flowing 35μL (typical chamber volume) of 1mg/mL solution of small unilamellar vesicles (SUVs, 50nm in diameter) in Membrane buffer (10mM Tris pH 7.4,2mM MgCl_2_, and 50mM KCl). The SUVs are prepared by extrusion and have a lipid composition (mol %) of 30% DOPE, 30% DOPC, 10% liver PI, 25% DOPS, 4.9% Brain PIP_2_, 0.1% BODIPY TMR-PIP_2_, and a supplement of 15% cholesterol. The chamber is incubated with the SUVs for 15min at RT and then washed with 5X volume of membrane buffer. 35μL of 2μM GTPγs-activated Cdc42 is then introduced in the chamber and incubated for 7min at 37°C, after which 35μL of 1μM ΔEVH1 N-WASP is introduced and incubated for 15 min at RT. Finally, the chamber is rinsed with *8*0μL of membrane buffer, loaded with actin and accessory proteins in polymerization buffer (10 mM Tris-HCl, 73mM KCl, 2mM MgCl_2_, 0.2mM EGTA, 2mM ATP, 1mM GTPγs, 2mM DTT, and 0.1mg/mL Glucose Oxidase, 0.018 mg/mL Catalase, and 0.2mM Glucose (anti-bleaching solution), as well as with 0.5mM creatine kinase and 5mM creatine phosphate (our ATP regenerating system), which sets the starting time of the experiment. Before visualization the chamber is sealed with grease.

*Solution of actin and accessory proteins*: The actin solution consists of a fixed amount of actin monomers (10μM), Arp2/3 complex (40nM), Profilin (2.5μM), CPs (50nM), and hCof (3μM). The solution is supplemented with variable amounts (0 − 75 nM) of the disassembly factors Cor1B and AIP1 at 1:1 molar ratio. Myosin II motors are added in the form of small filaments, roughly consisting of 50 myosin dimers *(32)*. The molar percentage of labeled actin monomers and myosin motors is 10 and 50 %, unless stated otherwise.

#### Microscopy techniques

Samples are imaged with a laser scanning confocal microscope (LSM *88*0, Zeiss Germany) controlled by ZEN software in fast Airyscan mode (Zeiss Germany) or with a 3i Mariana (Denver, CO) spinning disk confocal microscope, equipped with a Yokogawa W1 module and a Prime 95B sCMOS camera, and controlled by Slide Book software. For both systems imaging is performed using a 63X⁄1.4 NA Corr. M27 Oil immersion Plan-Apochromat objective in combination with an Immersol 518F immersion media (Refractive Index *RI*_immersion_ = 1.518, at 23°C, Carl Zeiss, Germany). Actin is excited at 4*88* nm and, PIP_2_ or myosin II motors, at 56*8*nm.

### Supplementary text

#### Image analysis and data quantification

Data quantification is performed using MATLAB (MathWorks, MA, USA), Python, Huygens Professional software package (Huygens; Scientific Volume Imaging, Hilversum, Netherlands), Origin (OriginLab Corp., MA, USA), and ImageJ software. Image analysis of the spinning disc confocal data is done on the raw images (of 512×512 pixels^2^) whereas the images acquired using LSM Airyscan mode (relevant only for the data presented in Fig. 2E,F are first deconvolved using Zen Black 2.1 (Version 13.0.0.0) software. The deconvolution is based on the Wiener filtering method using default settings.

The confocal images are corrected for photobleaching, and for lateral and axial drifts using a ‘registration’ function from the ANTs package *(12)*. We employ a Rigid transformation algorithm (which includes both rotation and translation) to correct the drift within the *x,y* plane and *z*-directions, passing Dense sampling parameters to achieve higher accuracy. Due to similar index of refraction of the objective immersion oil and our sample (true for both the SLB and the actin cortex, see ‘3D image reconstruction’ below), image correction for index refraction mismatch can be neglected *(13)*.

##### Analysis of actin cortex thickness dynamics, network turnover speed and myosin motors active transport

From the stabilized confocal images, we extract the mean intensity of the actin monomers and myosin motors in the bulk solution (above the actin cortex), which is subtracted to deduce the polymerized F-actin network and myosin II motors densities and their distributions across the cortex thickness, which we use in combination with image segmentation to characterize actin cortex thickness dynamics (spatio-temporal evolution and steady state properties), F-actin disassembly rate, actin network turnover velocity, F-actin/myosin II colocalization, and myosin II motors active transport within the cortex and their effect on actin cortex dynamics.

*Bulk actin monomers (G-actin) intensity* We extract the intensity distribution of the actin monomers localizing in the bulk solution by manually selecting a voxel volume of 10×10×10 pixels3, positioned ≥ 10 pixels above the visually inspected upper cortical network boundary. Via this we minimize the effect of the fluorescence emitted from the actin cortex on the resulting intensity distribution. Furthermore, by applying MIP on a stack of 20 images, taken below and above the voxel center, we also confirm that network debris are not present in the selected volume, which would lead to artifactually high intensity values. Then, we fit the actin monomer intensity voxel histogram to a Gaussian function, from which the mean intensity and standard deviation *σ* are extracted.

Next, we subtract the average monomer intensity value from which the intensity of the polymerized (F-actin) network is deduced. The images are then binarized. Morphological dilation using a structural element ball with a radius of 3 pixels is applied to the binarized image to compensate for small holes in the network. The result is then divided into connected components, and the largest component is defined as the actin cortex area. For image binarization we choose a threshold given by *xσ*, where *x* is a parameter which we fine-tune to locate the segmented *upper* cortex boundary at the *z*-position where the F-actin intensity (measured perpendicular to the SLB) drops to half its peak value (as illustrated in Fig. S6). If the distribution consists of more than one peak (e.g., as commonly seen during the transient growth phase) we refer to the peak as the closest to the cortex upper boundary. Note that the *x* used for precisely locating the segmented surface at the inner cortex edge can differ from the one used for the upper segmented surface.

*Cortex segmentation and thickness evaluation* For each *x,y* position along the cortex we locate the first (and last) *z*-plane (from above) with a finite (non-zero) intensity value, to generate the cortex, outer and inner, *z*_outer_ (*x,y,t*) and *z*_inner_ (*x,y,t*), segmented surfaces. In side-view representations it is given by a line that follows the topography of the cortex upper (or lower) surfaces (see Fig. S4B). Since the inner cortex edge is located at the SLB surface, we can alternatively extract it from the SLB normal intensity histogram, specifically, from the SLB peak intensity position, denoted as < *z*_membrane_ >_*x,y*_. The peak position accuracy in our experiments ranges between 10 − 15nm, as deduced from the standard error of the mean, SE. Both are extracted by fitting the surface-averaged, SLB normal intensity histogram to a Gaussian function.

From the above quantities we can extract the, local and surface-averaged, temporal cortex thickness, *h*(*x,y,t*) = *z*_outer_ (*x,y,t*) −< *z*_membrane_ >_*x,y*_ and < *h*(*t*) >_*x,y*_ =< *z*_outer_ (*t*) >_*x,y*_ −< *z*_membrane_ >_*x,y*_, respectively. In experiments where the membrane is not labeled the position of the inner cortex edge can be deduced from normal F-actin distribution and it corresponds to the place where the intensity is on average 1/4 the F-actin peak intensity value (see blue horizontal line in Fig. S6).

##### F-actin network disassembly rate τ^−1^ and turnover speed *v*_*turn*_, myosin II active directed motion and transport towards the actin filaments’ tip

The actin disassembly rate, **τ**^−1^, can be evaluated from the decay of the average fluorescence intensity of the F-actin density at the cortex’s outer edge over time, during the transient growth regime. The speed of network turnover *v*_turn_ is extracted from the slopes of kymograph plots generated from, *xz*/*yz*, cross-section image sequences. Similarly, the motion of large individual motors clusters is extracted from Maximal Intensity Projection (MIP), *xz*/*yz*, cross-section image sequences. The dynamics of the population of the smaller myosin II filaments that reside in the network is quantified from the changes of the myosin density peak position *z*_c_ (SE = ±30nm accuracy) and average intensity *I*(*z*_c_), extracted from the fit of the normal myosin density distribution to a Gaussian function.

##### 3D image reconstruction

For 3D image reconstruction we use the Huygens Professional 21.04 software package (Huygens; Scientific Volume Imaging, Hilversum, the Netherlands). The spinning disk confocal raw images (TIFF format) are first deconvolved. The default settings are based on the Classic Maximum Likelihood Estimation (CMLE) algorithm. For data deconvolution we find that n = 1.4*8* show the best results, in accord with the measured refractive index of dense cytoskeletal networks and lipid membranes *(14)*. This suggests that image correction for index refraction mismatch, between the objective immersion fluid and our sample can be neglected *(13)*, which is not the case for more dilute crosslinked actomyosin networks *(15)*.

##### Actin cortex growth and steady state dynamics are rate limited

This can be demonstrated by showing that the diffusion “velocity” of actin monomers (the most abundant protein in the soup) travelling from the cortex outer edge to the SLB (which corresponds to the farthest travelled distance) is significantly higher than the actin cortex growth rate or turnover velocity. We measured the diffusion constant of actin monomers in reconstituted branched networks 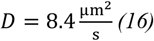, from which we can derive the monomers’ characteristic diffusion “velocity”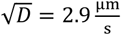. This value is two orders of magnitude faster compared to the measured network growth and turnover velocities, the latter being in the range of 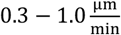. This suggests that even for very thick cortical networks (e.g., tens of microns), the time that it would take for the actin monomers to reach the SLB is just a few seconds, which would signify that actin cortex growth is not limited by diffusion. We can safely assume that this estimate is also valid for the rest of the proteins present in the actin solution, as well as for the Arp2/3 complex which is slightly larger in size. Although myosin II motors do not directly participate in actin cortex assembly, they do affect the rate of network assembly and the speed of actin network turnover. Similarly to the other proteins they reach the SLB very quickly (≪ 1 min) owing to their active directed transport capabilities, much faster than the characteristic rates of actin network growth and turnover, further supporting the fact that actin cortex dynamics are rate limited.

## Theoretical Model and Phenomenology

### Reaction-Diffusion-Advection model equations

Following the properties of the actin cortex dynamics observed and discussed in detail in the main text, we devise a qualitative minimal model that incorporates the basic kinetic and transport properties. The model accounts for the primary soluble components in one spatial dimension (the coordinate *z*, perpendicular to the surface of the suspended lipid bilayer (SLB), as follows:

- Polymerization of freely diffusing G-actin monomers (*m*(*z, t*)) into a rigid F-actin network (*p*(*z, t*)) at the SLB;
- Propagation of the F-actin network from the SLB surface into the bulk, where the speed depends on the conversion at the SLB;
- In addition to the actin monomers, freely diffusing disassembly agents (*c*_*f*_ (*z, t*)) to which in the main text we refer as [CA];
- Diffusing disassembly agents bind to F-actin, resulting in F-actin bound to disassembly agents (*c*_*b*_ (*z, t*)) that convert the F-actin (*p*(*z, t*)) into actin monomers (*m*(*z, t*)) and to freely diffusing disassembly agents (*c*_*f*_ (*z, t*));
- the total amount of material is conserved on the relevant time scales of the experiment: 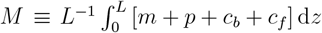, where *L* is the domain length and for convenience is taken as *L* = 1.

Consequently, the resulting model equations read:

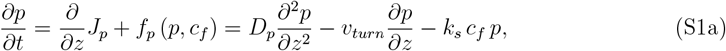

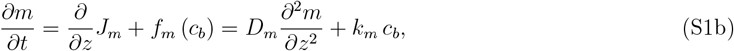

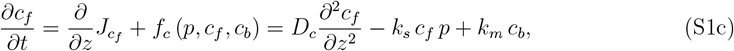

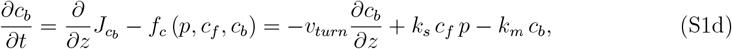

where the speed of the F-actin network is driven by the rate of actin polymerization at the SLB,

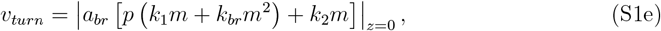

where different *k*_*i*_ are rate constants, and *a*_*br*_ is an arbitrary length scale converting the actin polymerization rate to speed. In (S1e), the leading polynomial terms containing *m, p* denote the incorporation of G-actin monomers into existing F-actin filaments: the first term denotes simple growth at existing barbed ends, while the second corresponds to the higher-order process of Arp2/3-induced branching. The last term in (S1e) describes the rate of spontaneous formation of new F-actin nuclei at the SLB from G-actin; for the sake of simplicity, we use only a linear dependence on *m*. To keep fidelity to realistic properties, as much as possible, we use *k*_1_, *k*_*br*_ *< k*_*m*_ *< k*_2_ and *D*_*p*_ ≪ *D*_*m*_ *< D*_*c*_. Based on the above assumptions, we use the following parameters: *D*_*m*_ = 8, *D*_*c*_ = 10, *D*_*p*_ = 10^*−*5^, *k*_*s*_ = 2, *k*_*m*_ = 0.2, *a*_*br*_ = 0.1, *k*_1_ = 0.0, *k*_*br*_ = 0.01, and *k*_2_ = 1. We note that the results qualitatively persist for *k*_1_, *k*_*br*_ ∈ [0, 0.1] while modifying the form of the steady states and the dynamics to reach it. Additionally, while the value of *D*_*p*_ ≪ 1 may account for minor branching in three dimensions, it is mostly used for numerical regularity.

For the boundary conditions at *z* = 0, that is, at the SLB, we employ conversion from G-actin to F-actin and no flux for *c*_*f*_,

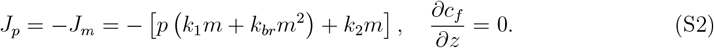

while at *z* = *L*, i.e., in the bulk, we set no-flux boundary conditions,

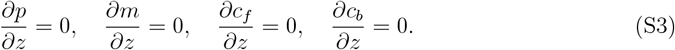

Finally, the initial conditions are

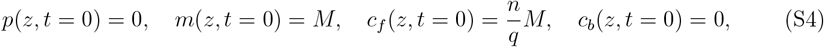

where for simplicity we use *q* = 8 and *n* values are taken as integers.

The simplification of system (S1) can be achieved via dimensionality and taking into account that *D*_*c*_ */D*_*m*_ ∼ 1 and *D*_*p*_ */D*_*m*_ ≪ 1, so that by introducing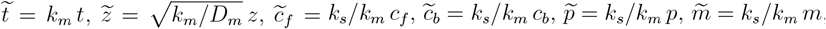, and after omitting the tildes, we obtain:

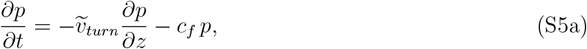

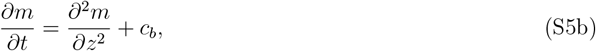

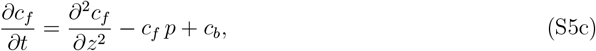

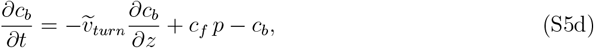

where

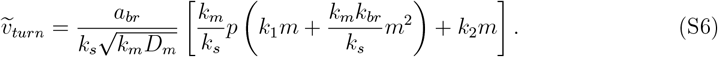

We see that *v*_*turn*_ is scaled with the model parameters that describe the processes that determine the supply of actin monomers: the disassembly rates of F-actin and the release of G-actin monomers *k*_*m*_, *k*_*s*_ and the diffusion *D*_*m*_ of these monomers to the SLB. Note also that *k <* 1, and that the functional forms for *v*_*turn*_ (Eq. S1e) and *J*_*m*_ (Eq. S2) do not change.

### Effective disassembly rate and the exponential cortical actin density profile at steady-state

The model described above contains two first-order processes that together give rise to the disassembly of newly formed F-actin (*p*) at the SLB: first there is the adsorption of free disassembly factors on the F-actin, at rate *k*_*s*_. This is then followed by the disassembly of the complex of F-actin decorated by disassembly factors (*c*_*b*_) at a rate *k*_*m*_, releasing actin monomers (*m*)and free disassembly factors (*c*_*f*_) into the solution.

We can neglect the differences in the spatial distributions, *∂*_*t*_ = *∂*_*z*_ = 0, of the different components, such that *c*_*f*_ is assumed to be uniform over the lengthscale of the cortical thickness, and approximate these two first-order sequential reactions by a single effective rate, as follows: the average time to pass through a sequential series of reactions that have constant rates is simply the sum of the average times (inverse of the rates)

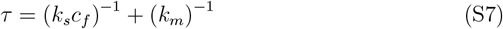

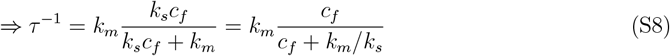

Note that the *c*_*f*_ in this equation is, in general, time-dependent. However, in the limit of a large reservoir, we can simply replace *c*_*f*_ by the concentration of free disassembly factors that were initially introduced into the system, such that *c*_*f*_ ∼ [*CA*], as appears in Eq.1 of the main text. If a significant amount of disassembly factors is removed from the solution by binding to F-actin, we have to include an additional proportionality factor (*c*_*f*_ = *α*[*CA*]), which can be parameterized by *K* in Eq. 1 (*K* = *k*_*m*_ */*(*αk*_*s*_)).

Finally, we can use the effective disassembly rate (Eq. S8), to further simplify and explain in simple terms the origin of the steady-state exponential density profile of the cortical actin (Fig. 2E in the main text). From the model formulated above, we have polymerized actin that forms at the SLB and is advected at speed *v*_*turn*_, and undergoes effective disassembly at rate *τ* ^*−*1^. A phenomenological advection-degradation can therefore be written for the density profile of the cortical actin if we are at steady-state conditions (i.e. when *v*_*turn*_ = *const*), of the form

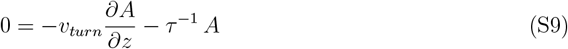

where *A* is the polymerized actin density field (effectively *p* + *c*_*b*_ from Eqs. S1). At steady-state, this equation has the solution of the form

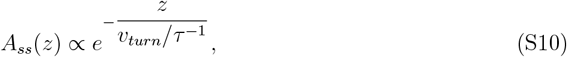

which shows that the steady-state thickness of the actin cortex is, therefore, expected to be given by the ratio

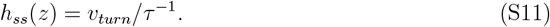

### Supplementary Figures

**Fig. S1:**
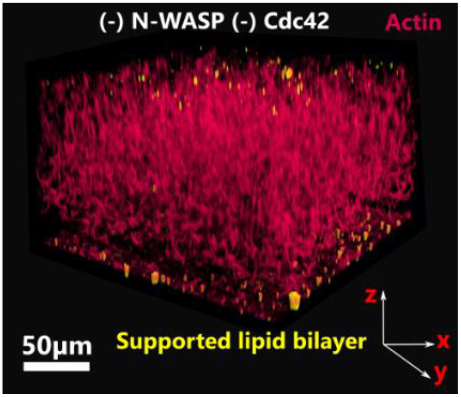
Actin cortices do not form in the absence of N-WASP and Cdc42. Shown is a 3D confocal image of system consisting of a SLB (yellow) coated flow chamber in which we have directly introduced a solution of pure components constituting of actin monomers (red, 10 mol% Alexa-Fluor 488 labeled) and actin accessory proteins (without CA and without myosin II motors), and without preincubating the SLB with GTPγs-activated Cdc42 and N-WASP. Under these conditions we can only detect actin filaments that spontaneously nucleate in the bulk solution that fills the whole chamber volume.

**Fig. S2:**
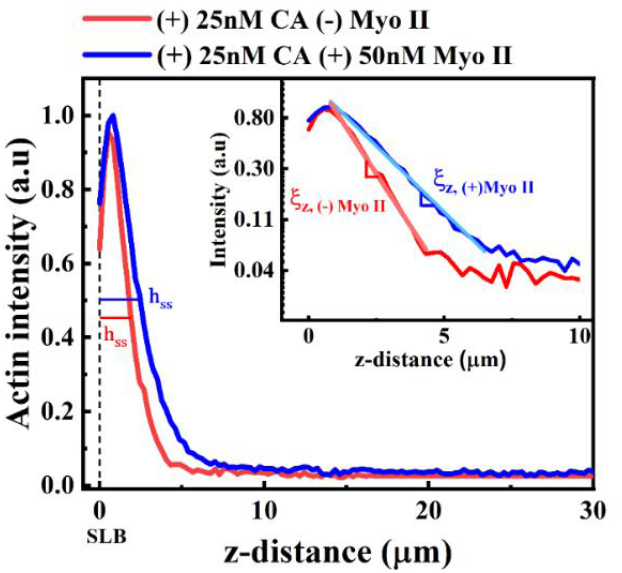
The actin density decays exponentially over the thickness in actin cortices exhibiting dynamic turnover. Shown are normalized steady state actin density (intensity) profiles measured perpendicular to the SLB for systems consisting of CA, with (blue line) or without (red line) myosin II motors. The steady state thickness *h*_*ss*_ corresponds to the distance from the SLB surface (located at *z* = 0, marked by the black dashed line) up to the *z* position which corresponds to the intensity at half maximum. Inset: Semi-logarithmic plots show that at steady state the actin density decays exponentially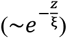, where ξ^-1^ is a characteristic decay length. This length is smaller in the presence of myosin II motors: ξ_(−)Myo II_ = 1.22 ± 0.03 μm < ξ_(+)Myo II_ = 1.93 ± 0.03 μm.

**Fig. S3:**
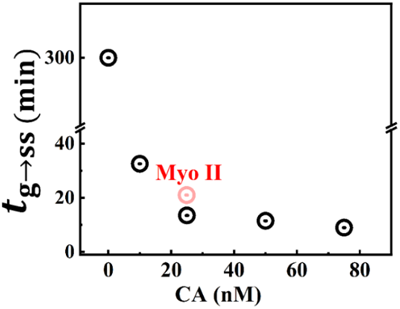
The transition time *t*_*g*→*ss*_ from transient growth to steady state regime shortens with the increase in [CA]. For [CA] = 0nM the system does not reach steady state, and it remains in the growth regime throughout the experimental time scale (typically 5 − 6h). The addition of 50 nM myosin II motors (red dot) to a system with CA = 25 nM delays the transition from growth to steady state regimes.

**Fig. S4:**
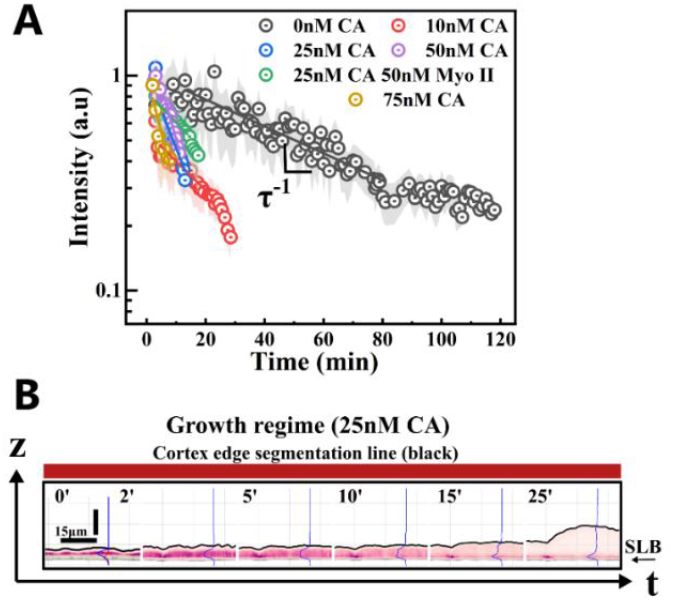
Extracting the network disassembly rate τ^−1^ from the decay of the fluorescence of the F-actin density at the cortex’s outer edge during the transient growth phase. **(A)** F-actin density measured at the cortex outer edge marked by the black segmentation lines in (B)) against time within the transient growth phase, for different experimental conditions (semi-log scale). Data corresponds to mean values (dots) while experimental errors (colored area) correspond to the standard deviation of the experimental data (*N* = 110). The semi-logarithmic plots show that the actin density decays exponentially with time, 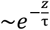, where **τ** is a characteristic decay time extracted from the slope. **(B)** Side-view sequential images of the time evolution of a growing actin cortex during its growth phase (25nM CA without myosin motors). The black lines mark the location of the cortex outer edge. Shown are also the density (intensity) distribution profiles of the F-actin (blue) normal to the SLB (thin black line marked by an arrow).

**Fig. S5:**
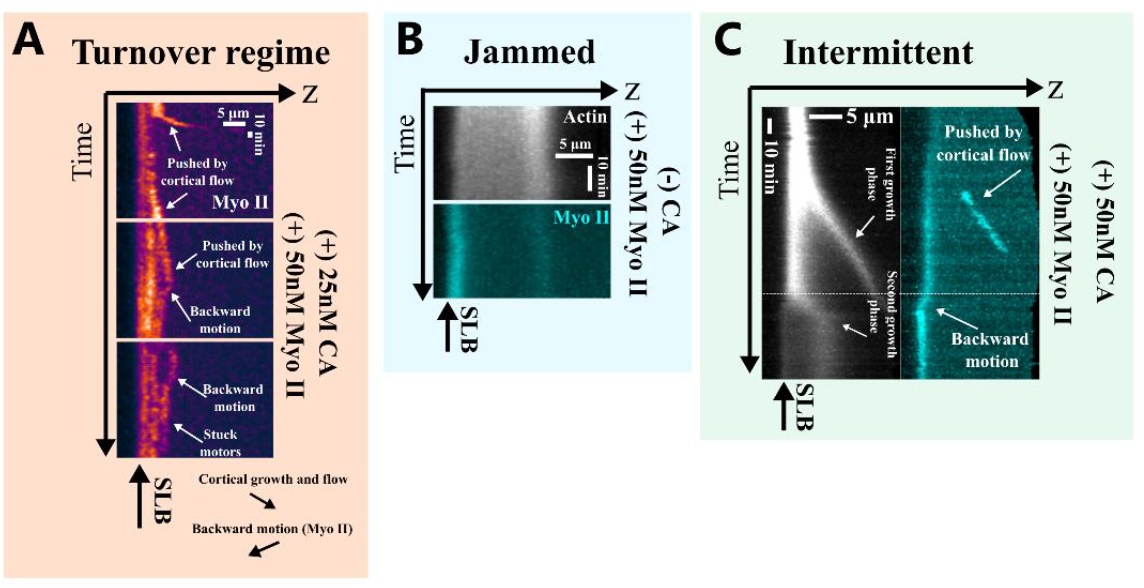
Actin polymerization dynamics and myosin active motion exhibit distinct behavior in different dynamic states. (**A**) Steady state dynamic turnover regime: Kymographs show trajectories of myosin II motor clusters. (Top) Two myosin clusters are pushed by actin treadmilling toward the cortex edge, where they are released into the solution and fuse. (Middle and bottom) Myosin clusters move bidirectionally: (i) pushed upward by actin flow and (ii) move backward toward the SLB, localizing at actin filament tips. Some motors become immobile (white arrows). (**B**) **Jammed regime**: (Top) Kymographs show actin and small myosin filaments in the jammed regime, where the network does not reach a turnover state. Myosin accumulates at the SLB surface, and actin and myosin densities inversely distribute across the cortex. (Bottom) **Intermittent regime**: Kymographs of actin and myosin II motors show intermittent growth and decay of the cortex. During the first actin burst (growth), a large myosin cluster moves with the flow. This is followed by a second burst (bottom arrow) and backward myosin motion towards the SLB, correlating with a decrease in actin network density.

**Fig. S6:**
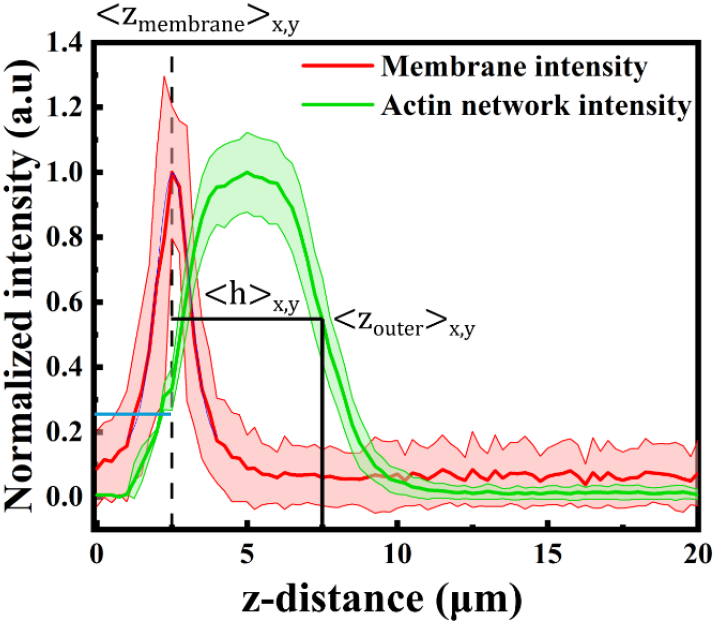
F-actin and SLB normalized intensity distribution profiles measured normal (perpendicular) to the glass coverslip used for extracting membrane peak position/inner actin cortex edge, actin cortex outer edge, and cortex thickness. Average normal intensity distribution profiles (thick middle lines) of the F-actin cortex (green) and the membrane (red) measured normal to the glass coverslip. The corresponding-colored area represents the standard deviation of the experimental data (*N* = 115 positions over the SLB/actin cortex surfaces). The SLB peak position < *z*_membrane_ >_*x,y*_ (black vertical dashed line) is extracted from the fit to a Gaussian function and has an accuracy of 10nm. The mean cortex thickness < *h* >_*x,y*_ = < *z*_outer_ >_*x,y*_ −< *z*_membrane_ >_*x,y*_ where < *z*_outer_ >_*x,y*_ defines the point where the F-actin intensity drops to half its peak value. In experiments where the membrane is not labeled, the position of the inner cortex boundary/edge corresponds to the location where the F-actin intensity is ∽1/4 the peak intensity from the left (blue horizontal line).

